# Behavioral Classification of Sequential Neural Activity Using Time Varying Recurrent Neural Networks

**DOI:** 10.1101/2023.05.10.540244

**Authors:** Yongxu Zhang, Catalin Mitelut, David J. Arpin, David Vaillancourt, Timothy Murphy, Shreya Saxena

## Abstract

Shifts in data distribution across time can strongly affect early classification of time-series data. When decoding behavior from neural activity, early detection of behavior may help in devising corrective neural stimulation before the onset of behavior. Recurrent Neural Networks (RNNs) are common models for sequence data. However, standard RNNs are not able to handle data with temporal distributional shifts to guarantee robust classification across time. To enable the network to utilize all temporal features of the neural input data, and to enhance the memory of an RNN, we propose a novel approach: RNNs with time-varying weights, here termed Time-Varying RNNs (TV-RNNs). These models are able to not only predict the class of the time-sequence correctly but also lead to accurate classification earlier in the sequence than standard RNNs. In this work, we focus on early sequential classification of brain-wide neural activity across time using TV-RNNs applied to a variety of neural data from mice and humans, as subjects perform motor tasks. Finally, we explore the contribution of different brain regions on behavior classification using SHapley Additive exPlanation (SHAP) value, and find that the somatosensory and premotor regions play a large role in behavioral classification.

## Introduction

Robust classification of behavior from multi-regional sequential neural data has attracted more and more attention^1,2^. Temporal neural activity can be classified sequentially in time, which has the potential for early detection of behavior. However, classifying the entire sequence and having a running estimate of the accuracy are often competing goals, especially with significant changes in the distribution of the data across time. Here, we investigate the accurate classification of behavior from neural time-series as early and reliable as possible. Specifically, we would like to predict the behavior before it happens, while having robust classification across time despite temporal distributional shifts in the data^3^.

Recurrent Neural Networks (RNNs) are designed for time-series data: they take in sequential inputs and predict the class of the sequence using recurrent hidden states that are able to retain a memory of previous inputs. However, standard RNNs are static in nature, and thus do not perform well on data with long time-series and shifting data statistics. Instead, they are good at accurately classifying temporal data at the end of the sequence. To help the network utilize all temporal features of the input and to enhance the memory of an RNN, here we propose a novel approach: RNNs with time-varying weights, termed Time-Varying RNNs (TV-RNNs). These models are able to not only predict the class of the sequence correctly, but also lead to accurate classification earlier in the sequence than standard RNNs. In this work, with TV-RNNs, we focus on the early sequential classification of brain-wide neural activity across time, as subjects perform a motor task (Fig 1A). Three different datasets are used: (1) **simulated data** with chirp signals to simulate distributional shifts in the data, (2) **widefield calcium imaging (WFCI)** records the neural activity across mouse dorsal cortex while subjects perform a behavior, here, ‘lever pull’ task, and (3) **functional magnetic resonance imaging (fMRI)** that records human whole-brain neural activity while patients with Parkinson’s Disease and healthy controls perform a ‘grip force’ task.

**Figure 1.**
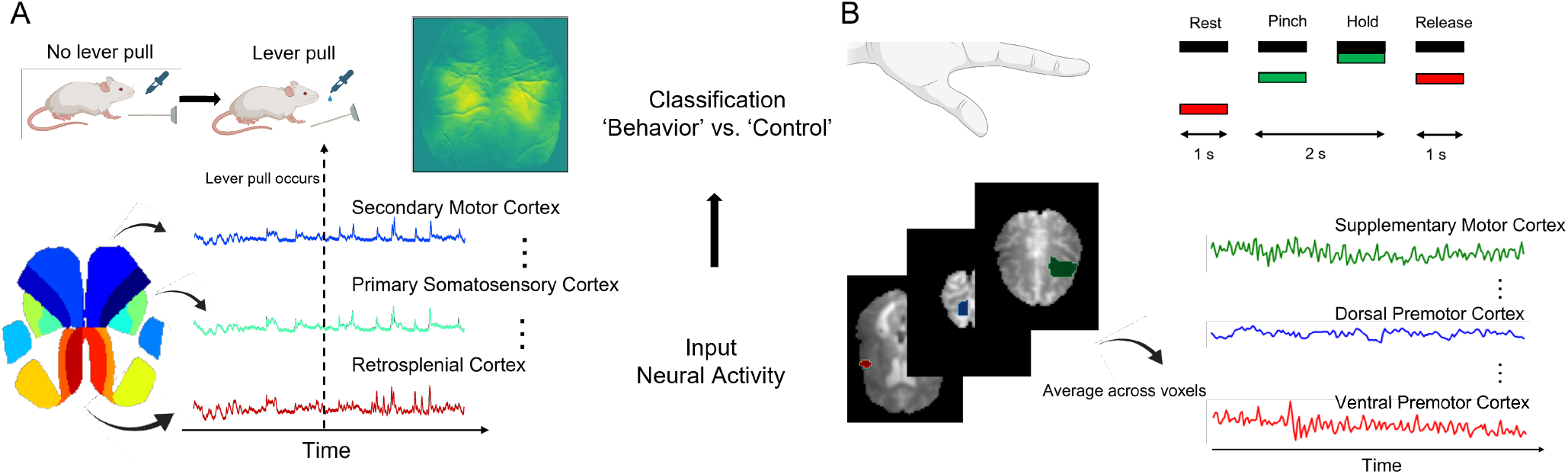
(A) In the WFCI dataset, mice were trained to pull a lever for water reward, while WFCI activity was recorded from multiple regions. (B) Neural activity of healthy and PD human subjects in a grip force task was recorded using fMRI.

We first examine the data distribution across time: the data distribution in the behavior trials changes across the trial, as quantified by visualizing the short-term Fourier transform (STFT). Due to the shifting data distribution, we hypothesized that TV-RNNs will perform early binary classification of behavior. As a comparison, we show the results while using two different loss functions to train standard RNNs using backpropagation-thru-time (BPTT): (a) the loss at the end of time sequence, which is a common strategy to train RNNs for classification, and (b) the loss at all time steps, which concentrates on not only the classification of the entire sequence but also at each time point in the sequence. As evaluation metrics, we consider the ‘temporal accuracy’ of an RNN, which quantifies the performance of the model at each timestep, and we aim to enable RNNs to perform classification as early and as accurately as possible. Moreover, we use SHapley Additive exPlanation (SHAP) values to visualize the effect of different regions on the classification of brain-wide data^4^. We find that the somatosensory and premotor regions play a large role in behavioral classification across both species. We show that (a) TV-RNNs outperform standard RNNs, (b) we are able to understand the classification mechanisms of TV-RNNs, and (c) we are able to accurately pinpoint the effect of different regions on behavioral decoding.

## Related Work

### Behavioral classification using neural activity

Previous research has focused on behavioral decoding by splitting data into multiple windows and independently applying separate classifiers to each window of data. For example, Soon et al. use a support vector machine (SVM) to decode the decision made by humans up to 10 seconds before awareness^1^; in our previous work, we have also succeeded in behavioral decoding using a sliding window approach^56^. However, the temporal information hidden in the time series data is not adequately utilized in these models because each classifier is independent. Thus, RNNs become a more effective model because they are able to process neural data while taking into account the temporal features.

### Sequential classification

Classification of sequence data has attracted extensive attention and can be applied in many areas, e.g., genomic analysis, information retrieval, and health informatics^7^. Information about the data is stored across the sequence; in our work, we consider neural data in which the features for predicting the behavior are not only distributed sequentially in time but also across different regions of the brain. By using and storing information across the sequence, RNNs are able to convert their representations across time to adapt to the task, and thus, they perform well in classifying sequential data^8^. For example, in^9^, the presence of heart disease can be detected by RNNs using electrocardiogram data, while in^10^, text sequences are classified by RNNs.

Research has been lagging on performing early classification on temporal data, although this is attractive since we would like to predict the class of a time series as soon as possible in order to be able to act on it. In Xing et al., the authors explored the minimal prediction length for neural networks to classify time-series data accurately^11^. In Mori et al., the authors optimized the early index and accuracy of a network at the same time^3^. Here, we use a time-varying approach to perform early and sequential classification of behavior using neural data.

### Time varying models

Models with time varying parameters are efficient in dealing with temporal tasks, they have unique parameters to utilize specific information at different times. For instance, switching linear dynamical systems (SLDS) and recurrent SLDS are designed to parse data sequences into coherent discrete units which help to capture distinct dynamics in different time periods of time-series data. Moreover, time varying parameter regression models have shown their utility in a range of applications such as economics^12,13^. Classification can also be improved by applying different parameters temporally, e.g., Yang et al., used multiple CNNs in parallel across time to classify time varying signals^14^, and Wang et al., show that time varying parameters outperform common machine learning approaches in the classification of EEG signals^15^. However, these methods lack connections between different parts of the models: temporal information that may be crucial for classification is not transmitted across time. On the contrary, the proposed TV-RNNs have explicit hidden states storing and transmitting temporal information across the entire sequence.

## Methods

In this section, we provide details of our model, termed TV-RNNs. We also introduce the metrics that we use to quantify the classification performance of RNNs and compute the importance of different brain regions. In addition, the experimental details of the datasets are also shown in this section.

### Standard Recurrent Neural Networks

We build a classification model with time-series neural data *x* ∈ ℝ^*R*×*T*^ from *R* different brain regions and *T* time points as the input, with the outputs as the different classes of behavior. Here, we implement a hidden recurrent layer with the *tanh* activation function, and a dense layer at the output with the *sigmoid* activation function *σ* to predict the binary class. Following are the equations of the RNN network.

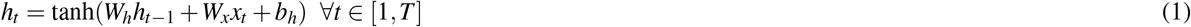

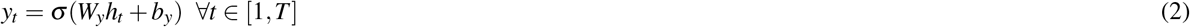

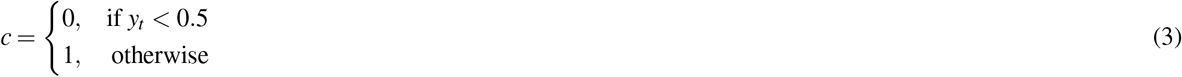

where *x*_*t*_ ∈ ℝ^*R*×1^ is the neural data from all *R* brain regions at time point *t, h*_*t*_ ∈ ℝ^*N*×1^ is the value for the *N* hidden units at time point *t, W*_*x*_ ∈ ℝ^*N*×*R*^ is the input weight matrix, *W*_*h*_ ∈ ℝ^*N*×*N*^ contains the recurrent weights for the hidden layer, and *W*_*y*_ ∈ ℝ^1×*N*^ represents the output weight matrix. *y*_*t*_ is the output of dense layer. Figure 2A shows the specific structure of the unfolded standard RNNs. We use backpropagation-thru-time (BPTT) to train the RNNs. We use two commonly used loss functions to train the standard RNNs: (a) the loss at the last output of RNNs (*y*_*T*_) in order to focus on the prediction of the entire sequence (written as ‘S1’); and (b) the loss at all time steps of RNNs sequence (∑_*t*_ *y*_*t*_), where we focus on not only the prediction at the end of the sequence, but also on the aggregate performance of the RNNs (written as ‘S2’). Note that only one set of weights need optimizing in both cases. The pseudo-codes are shown in Algorithms 1 and 2 respectively.

**Figure 2.**
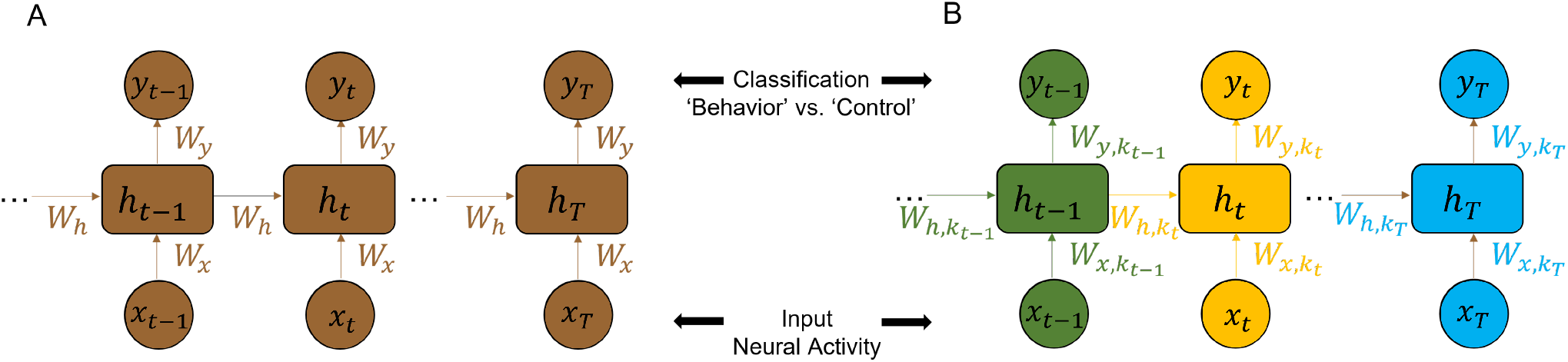
(A) standard RNN and (B) time varying RNN (TV-RNN) used for behavioral classification of neural activity from different brain regions.

#### Algorithm 1 RNN-S1

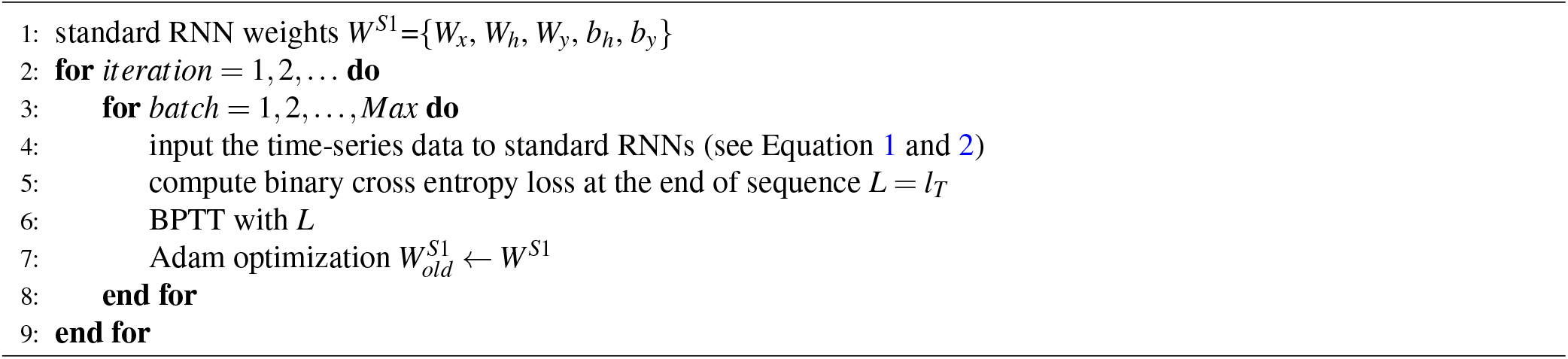

#### Algorithm 2 RNN-S2

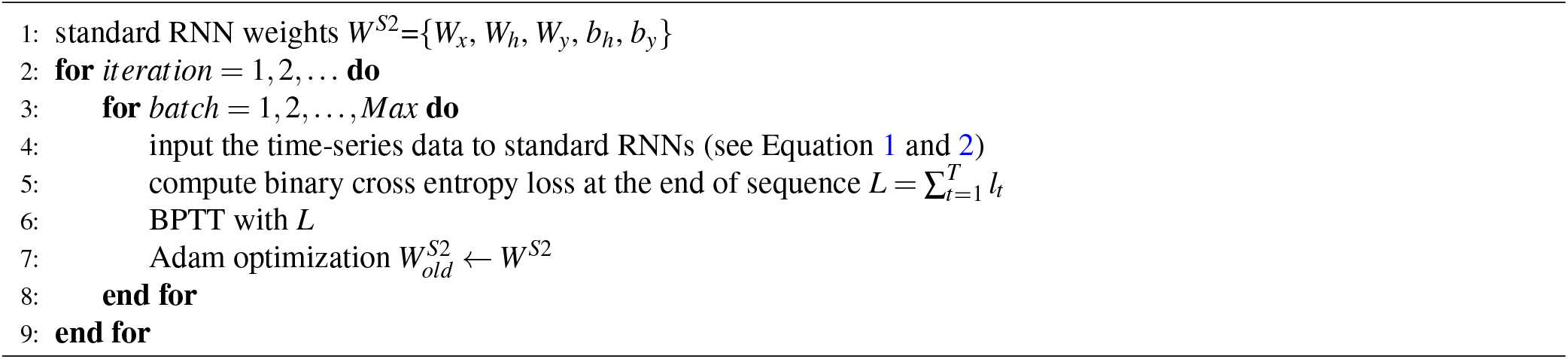

### Time Varying Recurrent Neural Networks

In order to capture the specific temporal features of the input, we design an RNN with time-varying weights including input weights 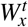, recurrent weights 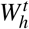, output weights 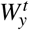, and bias 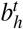, 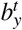.

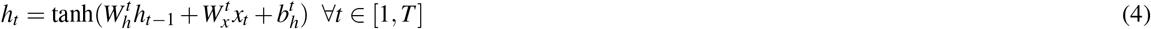

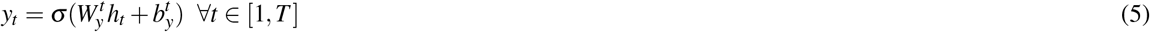

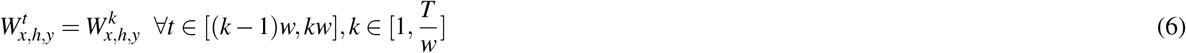

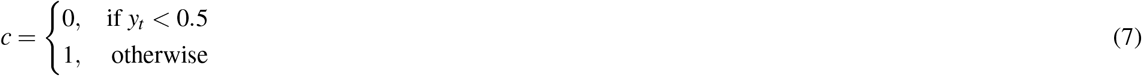

where *w* is the window size of RNNs. Specifically, the inputs in each time window *w* are fed into RNNs with one set of input weights 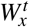, recurrent weights 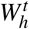, output weights 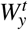, and bias 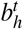, 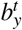. Thus, for the entire sequence of inputs, 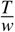 sets of weights are used. Fig 2B shows the specific structure of the unfolded TV-RNNs. In TV-RNNs, multiple sets of weights need to be trained. The optimization of all the TV RNN weights is performed simultaneously (end-to-end) with the same BPTT, i.e., in each batch training. The pseudo-code of training TV RNNs is shown in Algorithm 3. In all cases of standard RNNs and

#### Algorithm 3 Time-varying RNN

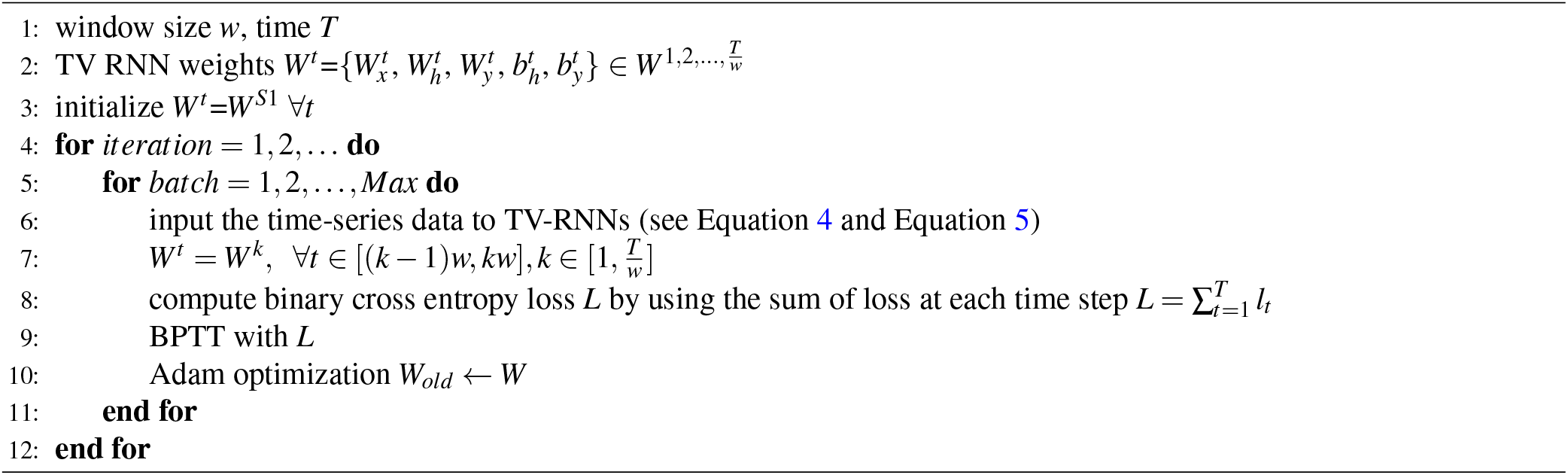

TV-RNNs, we chose the number of hidden units as 64, and we trained all networks for 1000 epochs using Adam at a learning rate of 0.0001. These hyperparameters were determined using cross-validation on a sample session of the dataset. We used Pytorch to train our models. We performed all tasks on HiPerGator Computational Supercomputer at the University of Florida, with NVIDIA A100 GPUs. The code is avaliable online https://github.com/saxenalab-neuro/TV_RNN.

### Accuracy Quantification

#### Temporal accuracy

In order to describe the performance of temporal decoding and exploring early classification, we use temporal accuracy *Accuracy*_*t*_, which depicts the classification accuracy at each time point *t*. We applied 5-fold cross-validation to all of our experiments. In the following, the data are split into 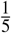 of test set in each fold, during training, the rest 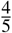 set is split into training set and validation set with validation rate of 0.2, the validation set is used to monitor over-fitting during training, and thus decide the hyperparameters, i.e., learning rate, training iterations, batch size, and the TV RNN window size *w*. All results are reported on test data. Accuracy here is defined as 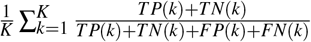, where *K* is the number of folds, here 5, *TP*(*k*) is true positives in the *k*^*th*^ fold, *TN* is true negatives, *FP* is false positives, and *FN* is false negatives.

#### Area under accuracy curve (AUAC)

We calculated the area under the accuracy curve in different time windows, above chance level. This quantifies the overall decoding ability of the classifier.

#### Earliest decoding time

We would like to classify the sequence as early and as accurately as possible. In order to explore the ability of RNNs in early classification, we introduce a metric, *earliest decoding time*, to measure early classification. This is the earliest time point after which we obtain consistent and significant decoding till behavior onset. Significance was determined using a one-tailed t-test at a significance level of *p <* 0.05 (after multiple hypothesis correction using the Benjamini-Hochberg procedure^16^). This represents the earliest time that the behavior can be reliably decoded.

### Effect of Brain Regions

The important features for decoding are stored in both the time domain and brain region domain of the data. In our previous work, we explored the occlusion method to quantify the importance of each region in both the time and brain region domain^6^. In this work, we employ SHapley Additive exPlanation (SHAP) value which is able to overcome the limitations of occlusion, because it considers the complete effect of a region on classification. SHAP value is firstly introduced by Lundberg and Lee in^4^. It interprets the effect of a given feature on the output of the model explained by computing Shapley values from coalitional game theory^17^. In binary classification, a positive value means the contribution of a given feature to predict a positive class, meanwhile, a negative value reflects the contribution to predicting a negative class. Therefore, we use the absolute SHAP value which is able to easily represent the contribution of the feature in binary classification. SHAP and importance are defined as follows:

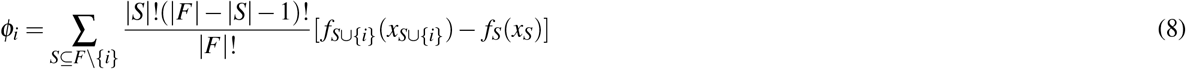

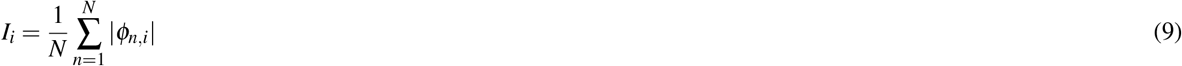

where *ϕ*_*i*_ is the SHAP value of feature *i, F* and *S* mean all the features and subsets of all the features, here, our features are in both time domain and brain region domain. Additionally, *f* represents our classifiers, i.e., standard RNNs and TV-RNNs, *I*_*i*_ indicates the importance of feature *i* which is an average of SHAP value of this feature across all trials. In this work, we employ GradientExplainer which is based on integrated gradient values to approximate the SHAP values. The integrated gradients are defined by Sundararajan et al., in^18^ as:

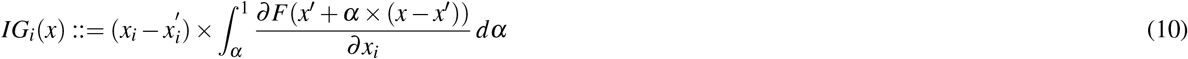

where *x* is the input used to explain our model, here the temporal neural activity in the test set, and *x*^*′*^ represents the baseline input. The integrated gradients are crucial to approximate the SHAP value.

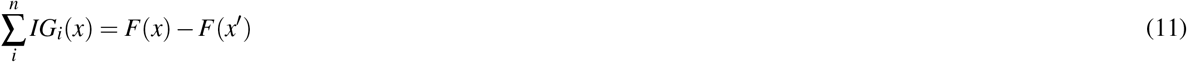

In our case, *F*(*x*^*′*^) is around 0.5 since we apply the *sigmoid* activation function in our output layer for both standard and TV-RNNs. Therefore, the difference between the outputs of subset with explained features and the subset without explained features in Eq.8 is able to be approximated by integrated gradients.

### Experimental Methods

#### Simulated Dataset

In order to uncover the effect of different features in both the region and time domains on behavioral classification and characterize the rationality of the method, we generated a simulated dataset with very clear features that we wish to recover using our methods. These features simulate the properties in our neural data that we would like to leverage for classification. The simulated behavior data consists of 10 chirp signals with 300 time points in each. Each of 10 signals is multiplied by a distinct coefficient selected within a range of (1, 4), and the amplitude linearly increases across time points in each signal. We then add Gaussian noise, and simulate 2000 trials. The simulated ‘control’ signals are shuffled ‘behavior’ signals across time.

#### Widefield Calcium Imaging (WFCI) Dataset

Widefield experiments record large-scale neural activity from the mouse dorsal cortex through WFCI. We analyze widefield neural activity while mice engage in a task. In the experiment, head-fixed water-deprived mice were trained to pull a lever and hold it at an angle (for *>* 100ms) in order to receive a water supplement. Rewarded lever pulls were identified online (using a lever analog signal), and a minimum 3 seconds lockout window was used to make sure the mice cannot get rewarded twice for less than 3 seconds. Widefield calcium imaging was recorded from the mouse dorsal cortex as previously described^19^. We identify the ‘behavior’ trials as trials that were tracked in real time to provide water reward, with the trial centered around the initiation of the lever pull behavior. As control trials, the time of the lever pull behavior was randomized to fall anywhere except a ±3 s window around an behavior, and we selected the same number of time points as the ‘behavior’ trials for the ‘control’ trials. Thus, the ‘behavior’ trials have a clear behavior initiated at the middle of the trial, unlike the ‘control’ trials. In order to further eliminate the influence of multiple instances of lever pulls occurring during a ‘behavior’ trial, we manually selected trials such that only one instance of lever pull is located at the middle of each ‘behavior’ trial. The neural activity is sampled at 30 time points per second, and each trial in this dataset contains 1800 time points (60 seconds). We spatially align the imaged neural activity with the Allen mouse brain coordinate framework^20^ using affine transformations, as previously performed in^21,22^. We then apply localized semi-nonnegative matrix factorization (LocaNMF)^22^on WFCI and take 16 components as identified by LocaNMF, which form our input signals, with each input dimension from one brain region. In this work, we focus on the signals around the lever pull, i.e., from 10 seconds before lever pull to 0 second after lever pull, because ‘behavior’ trials and ‘control’ trials are easier to be classified during these periods^56^. To increase trial counts, we use data pooled across all sessions of each mouse. Additionally, we show the quantification of classification performance using per-session data in sessions with greater than 39 trials to maintain a large trial count.

#### Functional Magnetic Resonance Imaging (fMRI) Dataset

An fMRI force production paradigm was used to assess differences in brain activity between patients with Parkinson’s Disease and healthy age-matched controls. Participants were required to perform a grip force task which consists of pinching the force transducer for 2 seconds, then releasing for 1 second, with visual displays presented during the task. Participants were asked to rest for 30 seconds before they start a grip force task, then perform the task for 30 seconds. They repeat this alternating rest-task procedure 4 times. Details are included in^23^. In our work, we use this dataset to test the ability of TV-RNNs to be applied to a large variety of neural data. We align the fMRI data to the Human Motor Area Template (HMAT) developed in^24^, and average the fMRI signals across voxels in each of 12 brain regions as shown in Fig 1B. The fMRI signal during subjects performing the grip force task is regarded as ‘behavior’, with the rest procedure being ‘control’, therefore, this is a binary classification task as well. Each trial has 12 time points representing 30 seconds in real time. In each group (patients or healthy control), we use RNNs to classify all ‘behavior’ and ‘control’ trials, i.e., grip force versus rest. We also compare the classification performance between Parkinson’s Disease patients and healthy controls.

## Results

We first analyze the data distribution of the datasets described above, and show that TV-RNNs outperform standard RNNs for them. We then examine the mechanisms of outputs of the RNNs, as well as the effect of different regions on the classification accuracy.

### Data distribution

We firstly show example trials of simulated behavior and simulated control signal in Fig 3A. In order to visualize the shifts in data distribution, we plot the STFT magnitude in Fig 3B. This reveals the data shifting over time from low frequency and low amplitude to high frequency and high amplitude. We also visualize the data distribution of the WFCI dataset (Fig 3C) and the fMRI dataset (Fig 3D-F). The STFT magnitude illustrate a shifts of data distribution existing in the WFCI data at around 5 seconds before the lever pull. However, the data distribution of the fMRI data is relatively static because the data has similar frequency and magnitude across time.

**Figure 3.**
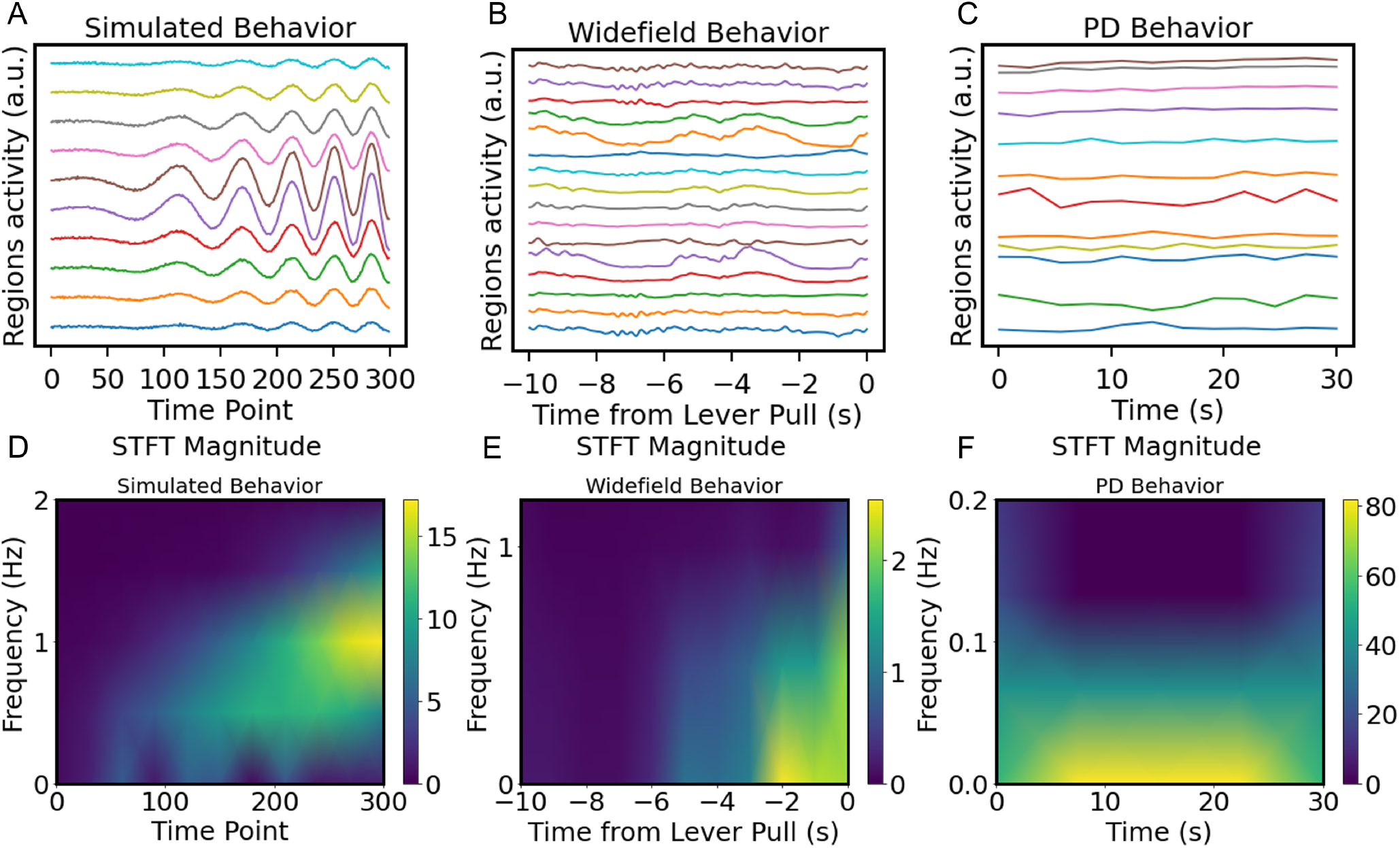
Plot of an example (A) simulated ’behavior’ trial, (B) WFCI ’behavior’ trial, (C) fMRI ’behavior’ trial. Short Time Fourier Transform (STFT) magnitude of (D) simulated behavior signal, (E) WFCI dataset and (F) fMRI dataset.

### Classification Accuracy Using RNNs

We compare the temporal classification accuracy of a simulated dataset between standard RNNs and TV-RNNs. We first use a common strategy: BPTT with a binary cross-entropy loss using the output at the end of the sequence (RNN-S1). Fig 4 (blue curve) shows that the temporal classification accuracy using this strategy only starts to increase above chance level after 150 time points. This trend matches the signal statistics in Fig 3D: after 150 time points, the simulated behavior signal has a higher frequency and magnitude. The alternative strategy to train Standard RNNs (RNN-S2), BPTT with the sum of the binary cross-entropy loss over time, leads to a low final accuracy (Fig 4, red curve). Consequently, a single set of weights in the standard RNNs does not seem to be able to guarantee early and accurate classification. On the other hand, TV-RNNs (Fig 4, green curve) can not only predict the class of the sequence early in the sequence, but maintain a high classification accuracy throughout the trial.

**Figure 4.**
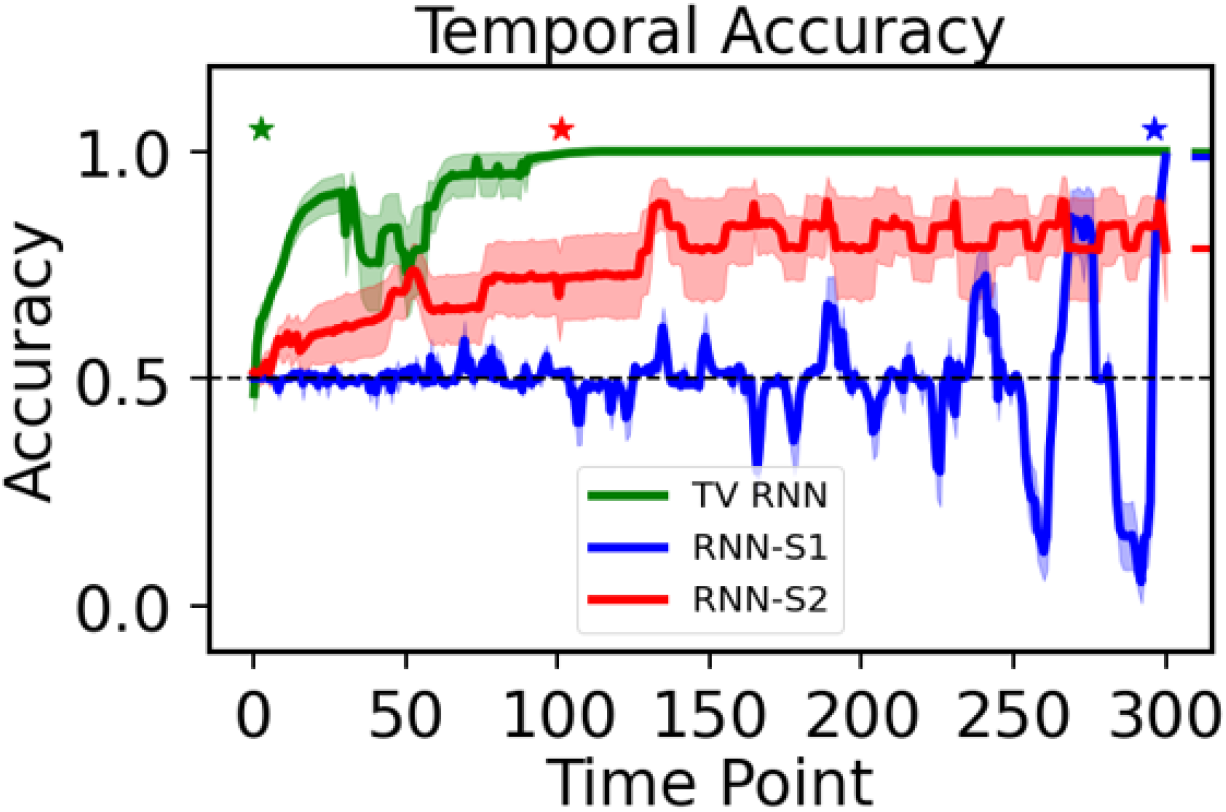
Temporal classification accuracy curve of standard RNNs and TV-RNNs using simulated data. The stars on top represent the earliest decoding time for each model (see Methods), and the bars on the right side reflect the final classification accuracy of the sequence. Note that chance accuracy level is 0.5 for both datasets.

We train the standard RNNs and TV-RNNs to classify the WFCI dataset of 300 time points, i.e., from 10 seconds before the behavior (lever pull) to the time that the behavior happens (see Methods), in order to quantify the earliest behavioral decoding time and the temporal performance of decoding with real data. We use an example session of one mouse with a large number of trials (here, 378 trials) to optimize the window length *w* between the values of 6 and 100, while computing the AUAC and earliest decoding time as metrics of interest (Fig 5A). Since *w* = 30 has the largest AUAC and the earliest decoding time, we set the TV-RNN *w* to 30, which implies that every 30 time points (1 second) will lead to a switch in the weights. Fig 5B shows the temporal classification accuracy with combined trials (2447 ’behavior’ trials). We see that the time around lever pull has the highest accuracy in both standard RNNs with S1 training strategy and TV-RNNs. Using standard RNNs, the behavior can be classified significantly above chance up to around several seconds prior to the lever pull, i.e., around 2 seconds in S1 and around 3 seconds in S2. This also illustrates that S2 performs better than S1 in early classification but worse in final classification, i.e., around 0.85 in S1 and 0.62 in S2. We also show that TV-RNNs significantly outperform standard RNNs in most time points in

**Figure 5.**
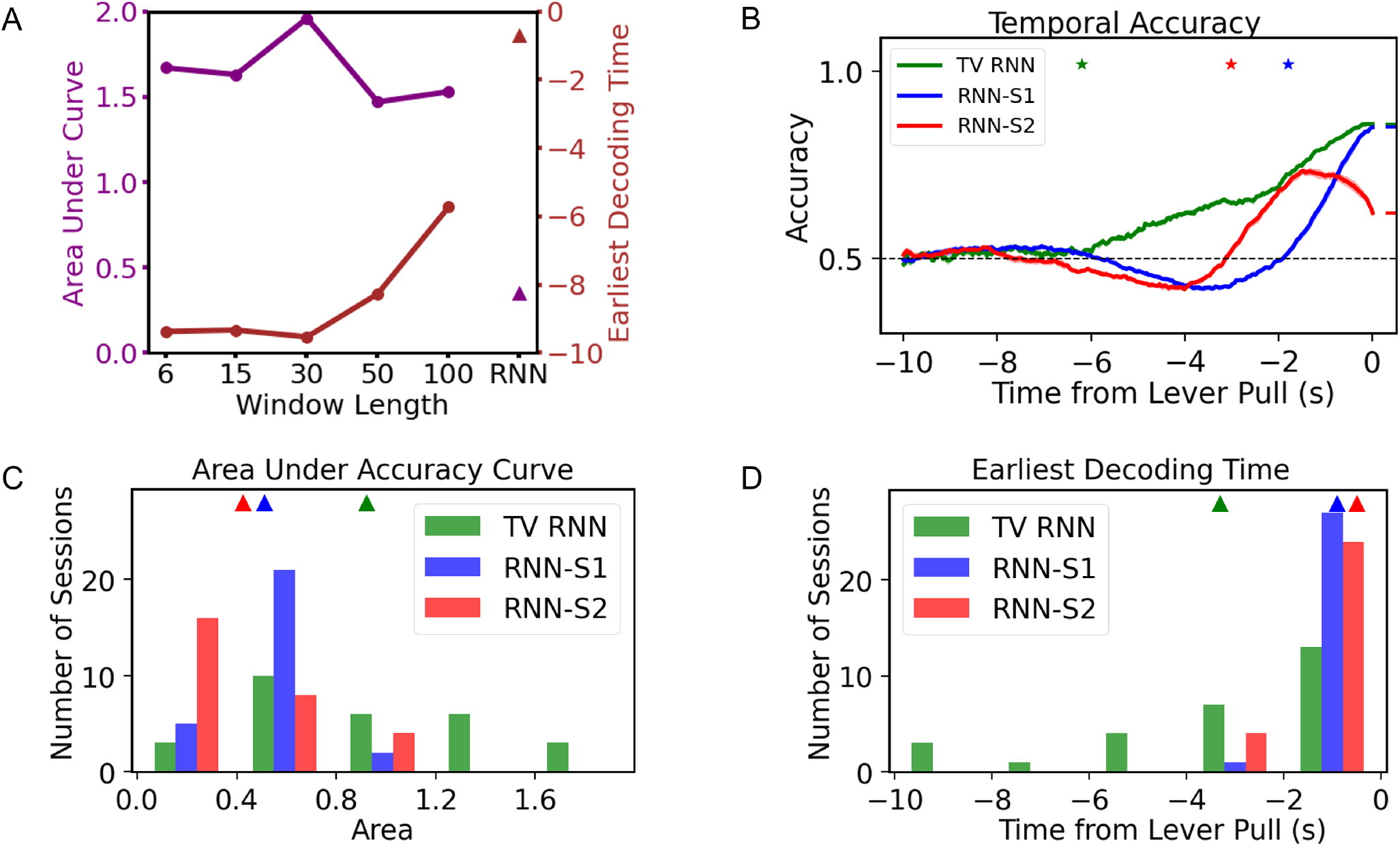
(A) Determining the window size *w* of TV-RNN: area under curve and earliest decoding time (see Methods) while varying *w* from 6 to 30; triangles represent standard RNN. (B) Temporal accuracy of standard RNNs with two training strategies and TV-RNNs, the stars depict the earliest decoding time with the height representing the sequential classification accuracy. (C) Histogram of the area under accuracy curve using standard RNNs and TV-RNNs for all sessions of mouse. (D) Histogram of the earliest decoding time using standard RNNs and TV-RNNs for all sessions of mouse.

Fig 5B. Importantly, the earliest decoding time of TV-RNNs can reach around 6 seconds before the lever pull, and the final classification accuracy is around 0.86. We also show the temporal accuracy curves of standard RNNs and TV-RNNs for another 5 mice with combined trials in S1. The TV-RNNs outperform standard RNNs in most time points as well, and all TV-RNNs have the earliest decoding time reach around 6 to 8 seconds before the lever pull. Next, we evaluate the session-by-session accuracy of the TV-RNNs. We first compute the AUAC in Fig 5C for selected sessions (with #*trials* ≥ 40). We consistently see that TV-RNNs achieve a higher accuracy than standard RNNs, presumably because TV-RNNs explicitly take into account more temporal structure with time-varying weights in the model, and allow for a monotonically increasing classification accuracy. Finally, we show the earliest decoding time in Fig 5D, where we see that TV-RNNs outperform standard RNNs in most sessions of the example mouse.

### Classification Mechanisms

In order to understand how the exact output of RNNs changes with shifting data distributions, we visualize the trained network activity succinctly: we plot the output of the networks (*y*(*t*)) in Fig 6C and D. The RNN output trajectories (Fig 6C) starts to diverge between the two classes at an early time, and at around 2 seconds before the behavior, the two trajectories from the two classes start to diverge quickly. Thus, the evidence for decision making between the two classes does not exist in the output nodes until close to the final time step *T*, at which point the information moves from the memory to the output nodes and the classification is performed. On the contrary, in Fig 6D, the TV-RNNs output trajectories start diverging at the beginning and towards the decision with accumulation of evidence^25^, the ‘behavior’ and ‘control’ trajectories start to diverge at around 5 seconds before the behavior. We also plot the same output trajectories for the simulated data. The standard RNN-S1 has overlapping trajectories between ‘behavior’ and ‘control’ at early times, and they cannot diverge well even near the end of the trial. The TV-RNNs output trajectories for simulated data have a similar tendency as the experimental dataset: they also diverge at the beginning and towards the decision with accumulation of evidence. Therefore, the TV-RNNs seem to utilize the temporal features in the data to accumulate evidence in order to make a decision. This is the reason why TV-RNNs not only outperform standard RNNs in Fig 4 and Fig 5B, but also why TV-RNNs are able to achieve monotonically increasing decoding.

**Figure 6.**
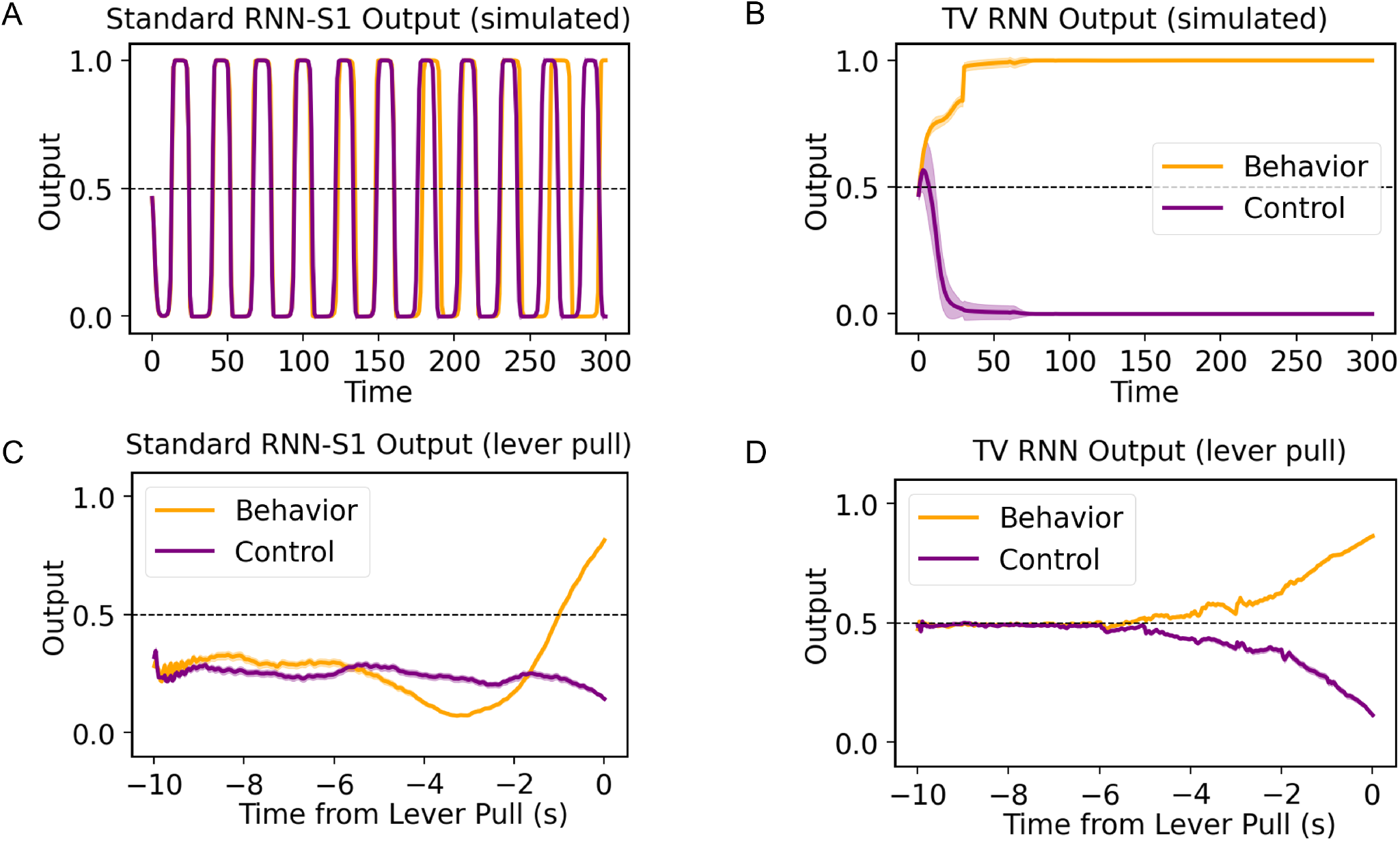
(A) Output trajectories of standard RNNs (average across trials), in the simulated data. The shaded region provides the standard deviation. (B) Similarly, the output trajectories of TV-RNNs. (C, D) Output trajectories for WFCI data: (C) standard RNNs; (D) TV-RNNs.

### Analysis of Weights for the WFCI dataset

In order to understand why TV-RNNs are more efficient at classification, and to analyze the difference between TV-RNNs and standard RNNs, we compare the learnt time-varying weights across the trial. We calculate the euclidean distance between the time-varying weights, i.e., 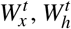 and 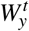, at different times in Fig 7A, B, and C. Note that the color-map illustrates the euclidean distance, with brighter colors representing a larger difference. We see that the weights are very different closer to the behavior. Furthermore, the recurrent weights 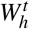 in Fig 7A shows more changes than the other weights, which may be necessary here to exploit the dynamic nature of the temporal features. Moreover, the sharpest changes in the input and recurrent weights are at around 3 seconds before the behavior, which matches the changes in the signal statistics in the STFT (Fig 3E).

**Figure 7.**
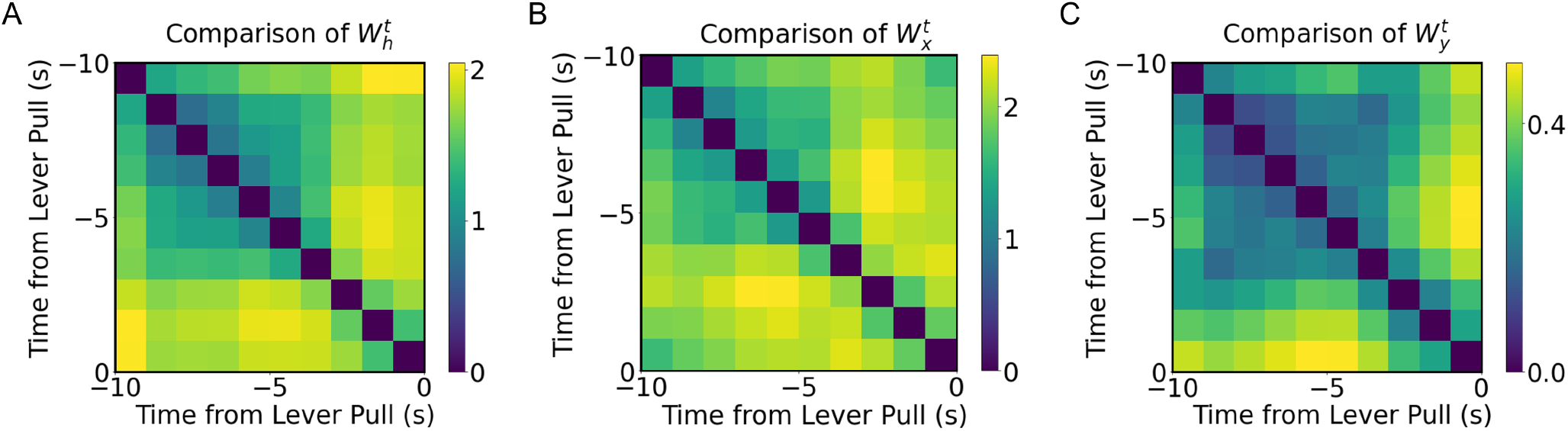
(A) Euclidean distance between 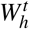 of TV-RNN at different time. (B) Euclidean distance between 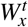 of TV-RNN at different time. (C) Euclidean distance between 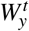 of TV-RNN at different time.

The output weights 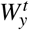 do not have a large variation, which reveals that the divergence between the trajectories of the two classes already exists in RNN layers. Consequently, the output weights can distinguish two classes without many changes.

### Comparison between TV-RNNs and other classifiers for the WFCI dataset

We compare the temporal classification performance of WFCI dataset between TV-RNNs and other classifiers. We first train 10 standard RNNs independently with each taking 30 time points (1 second) of the entire 300 time points (10 seconds before the lever pull). All the 10 standard RNNs have initialized hidden states of zeros. Note, these are different from TV-RNNs in which all sets of weights are dependent because of the continuous hidden states. Additionally, we train the 10 standard RNNs by using strategies RNN-S1 and RNN-S2 seperately as well. S2 shows the temporal classification accuracy of TV-RNNs and independent standard RNNs with combined trials for all 6 mice. We see that independent standard RNNs with S1 training strategy (purple curves) have large oscillations. This time, the earliest decoding time of all the mice is around 1 second before the lever pull, and the final accuracy of them is lower than the final accuracy of TV-RNNs. This matches the finding of standard RNNs with S1 training strategy in Fig. 5B, i.e., standard RNNs trained by using the loss at the end of the sequence are not able to classify the sequence accurately at early time. In contrast, independent standard RNNs with S2 training strategy (orange curves) can classify accurately at early time but not as accurately as TV-RNNs at final time. We then compute the AUAC of TV-RNNs and independent standard RNNs in S4, the TV-RNNs outperform independent standard RNNs at most time points, it illustrates that TV-RNNs do better not only because they have more weights than standard RNNs.

We then compare the TV-RNNs with support vector machine (SVM). We built SVMs using as input 1-s-wide windows (30 time points) of data. The input to each SVM classifier was a 2D array, i.e., [#*trials*,#*timepoints*\*#*components*]^5^. The classification accuracy of each window is reported as the temporal classification accuracy of the last time point in this window. We train SVMs with linear kernel and radial basis function kernel (RBF kernel). Here we only show the results of SVMs with RBF kernel in S3 and S4 because of its better performance than linear kernel. The SVMs outperform TV-RNNs in earliest decoding time but have lower final accuracy than TV-RNNs. The AUAC of TV-RNNs and SVMs are comparable. Indeed, sliding SVMs have 300 classifiers with each of them trained for 1 time point specifically, thus, it is fitter to each time point, especially at the early time points which have rare behavioral relevant information. Additionally, the classification accuracy after 2 seconds from the earliest decoding time is only slightly above the chance level. Furthermore, TV-RNNs outperforming sliding SVMs at final accuracy reveals that TV-RNNs can accumulate information for classification which multiple independent classifiers cannot.

### Performance on fMRI data

In order to test the performance of TV-RNNs on the fMRI dataset, we apply the same methods towards data from PD patients and healthy controls (see Methods). After exploring the best *w* of the TV-RNNs using the same methodology as for the WFCI dataset, we set *w* to 2 timepoints with this fMRI dataset. We train the standard RNNs and TV-RNNs by using the neural signal recorded from PD patients; the temporal accuracy curves are shown in Fig 8A. Here, TV-RNNs are able to not only classify the sequence accurately at the end but also achieve early classification, at around 2.5 seconds after the beginning of the sequence. Moreover, the temporal accuracy curves of healthy subjects in Fig 8B illustrate that TV-RNNs outperform standard RNNs and can achieve accurate classification at extremely early times, i.e., at the beginning of the sequence. Here, unlike for the simulated and WFCI datasets, the standard RNN-S2 is able to predict correctly at a relatively early time and keep high final accuracy in both PD patients and healthy control. One possible reason is that the subject starts applying a grip force from the beginning of the task, and the length of the sequence is shorter (only 12 time points), and thus standard RNNs are able to memorize most of the previous input. Another potential reason is that, as is evident in the STFT magnitude (Fig 3F), the data statistics are stable across time unlike the distributional shifts present in widefield datasets. Lastly, the classification accuracy is higher in healthy subjects than in PD patients, which can give us the insight that healthy subjects perform the finger moving task better than PD patients, who usually have a tremor in one hand.

**Figure 8.**
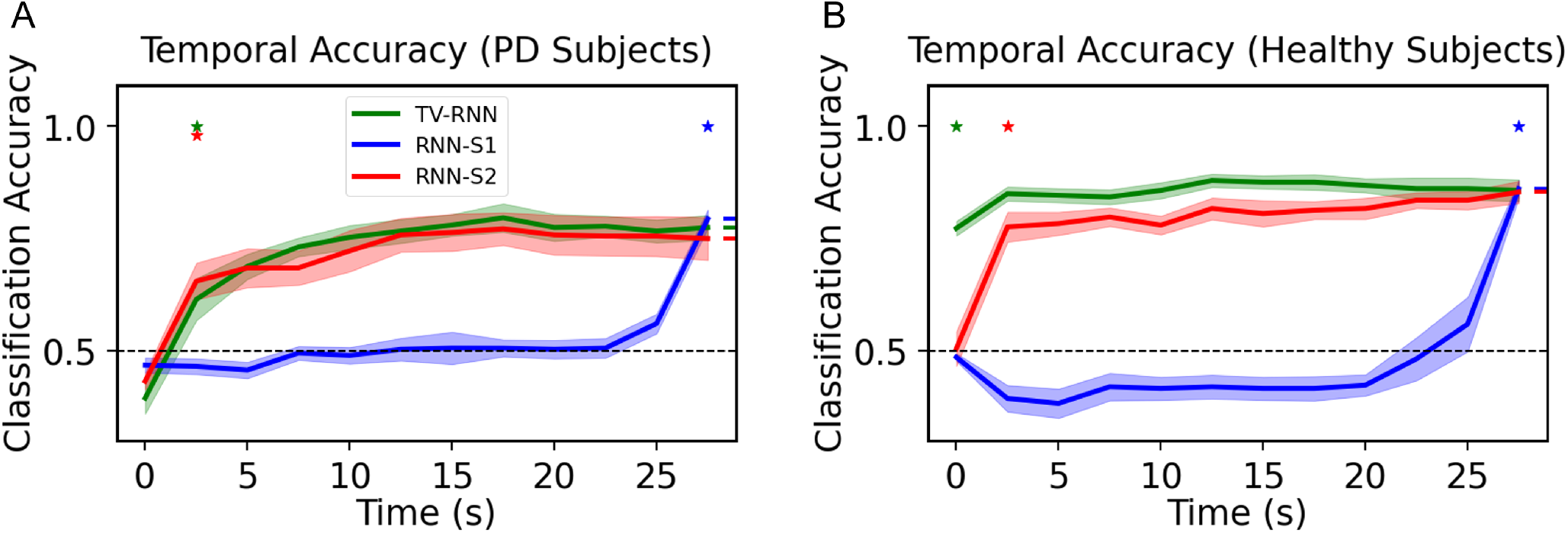
(A) Temporal accuracy of behavioral classification between ’force’ and ’rest’ for PD patients; (B) Temporal accuracy of behavioral classification between ‘force’ and ‘rest’ for healthy control.

TV-RNNs can be regarded as the best choice in early and accurate classification because of their best overall performance, as they can achieve both early classification and accurate sequential classification.

### Quantifying the contribution of different regions

#### Simulated Data

We compute the importance matrix of simulated data by calculating the average absolute SHAP value of each feature across all trials in the test set of the datasets (see Methods for details). The matrix in Fig 9A recovers the structure built into the trials, i.e., the consistent presence of the peaks in different dimensions at sequential time windows determines which class is the output. In Fig 9B, the importance is almost the same across regions. Only a few time points at the end of the sequence show relatively higher importance. However, the importance value is around 10^−6^. This is because that TV-RNN uses all the features at each region and time point, and all the features play a similar role in decoding. This points to a robust encoding of the behavior in the feature set.

**Figure 9.**
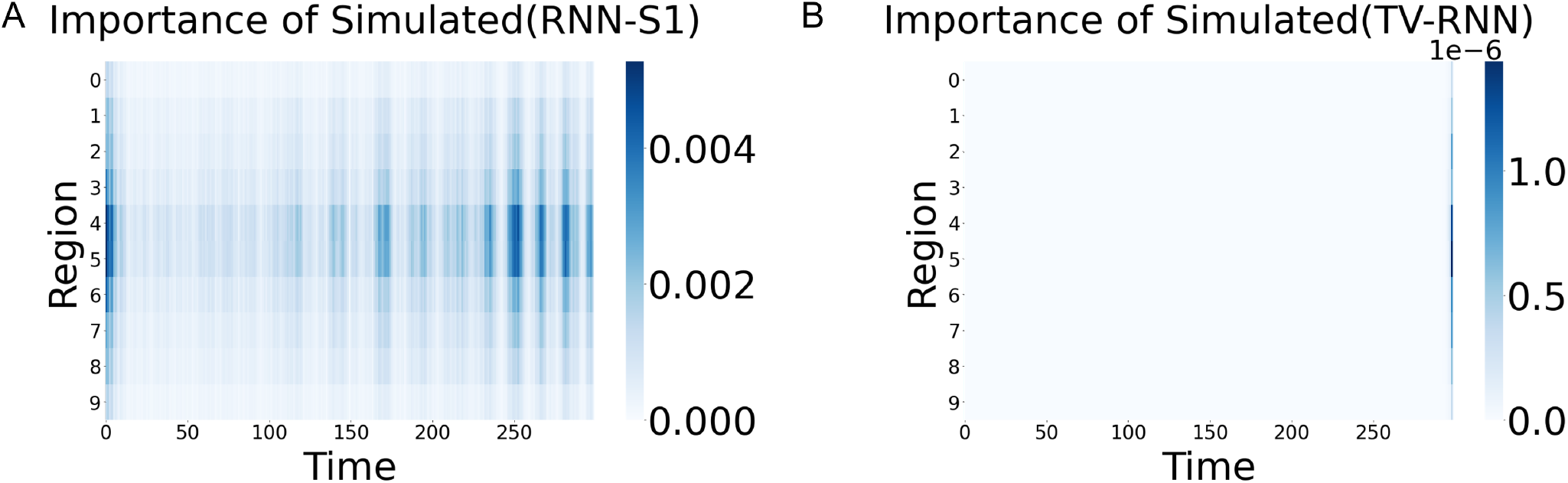
(A) Importance matrix of simulated data with standard RNN; (B) Importance matrix of simulated data with TV-RNN.

#### WFCI Dataset

Next, we examine the contribution of brain regions on ‘lever pull’ behavior with combined trials (the same mouse as Fig 5B). We compute the importance from SHAP value by using all the temporal outputs of TV-RNNs. Here we show five of them, i.e., 8 seconds, 6 seconds, 4 seconds, 2 seconds, and 0 second before the behavior respectively in S5, which means the contribution of brain regions on classification at 8 seconds, 6 seconds, 4 seconds, 2 seconds, and 0 second before the behavior respectively. These five importance matrix reflect the characteristic of RNNs, the input is forwarded into RNNs temporally, and the features of the input fed into the models after the target output time do not influence the target output. In the brain region domain, the left somatosensory upper limb region and the right somatosensory lower limb region show more importance than other regions across time, these two regions are considered to receive feedback from the right paw of the mice as it pulls the lever and to keep their body in balance. The regions with high importance are not always the same among different mice, showing subject-to-subject variability for mice performing the same task; the right somatosensory barrel field region, the motor regions, and the left somatosensory lower limb regions also show importance for other mice.

In Fig 10, we show the temporal importance of different brain regions while classifying the behavior of the example mouse performing the same self-initiated behavior by using TV-RNNs (Fig 10A, B, and C) and standard RNNs (not shown). According to the SHAP value at the end of the sequence, which is considered the final classification output, the somatosensory regions have more importance than other regions. In addition, the motor area in dorsal part is also important in one example session (Fig 10B). The left regions show more importance than the right regions (Fig 10A and C,), since all the mice used their right paw to pull the lever. In addition, the mice used their left paw to keep their body in balance when they pull the lever, so we can also see importance appearing in the right regions. Moreover, we found that when the mice used their paw to pull the lever, their lower limbs also moved, which may explain the finding that the somatosensory lower limb region is shown as important. The importance matrix of standard RNNs indicate similar important regions as the TV-RNNs case but less variable across time. This matches the mechanism of standard RNN and TV-RNNs.

**Figure 10.**
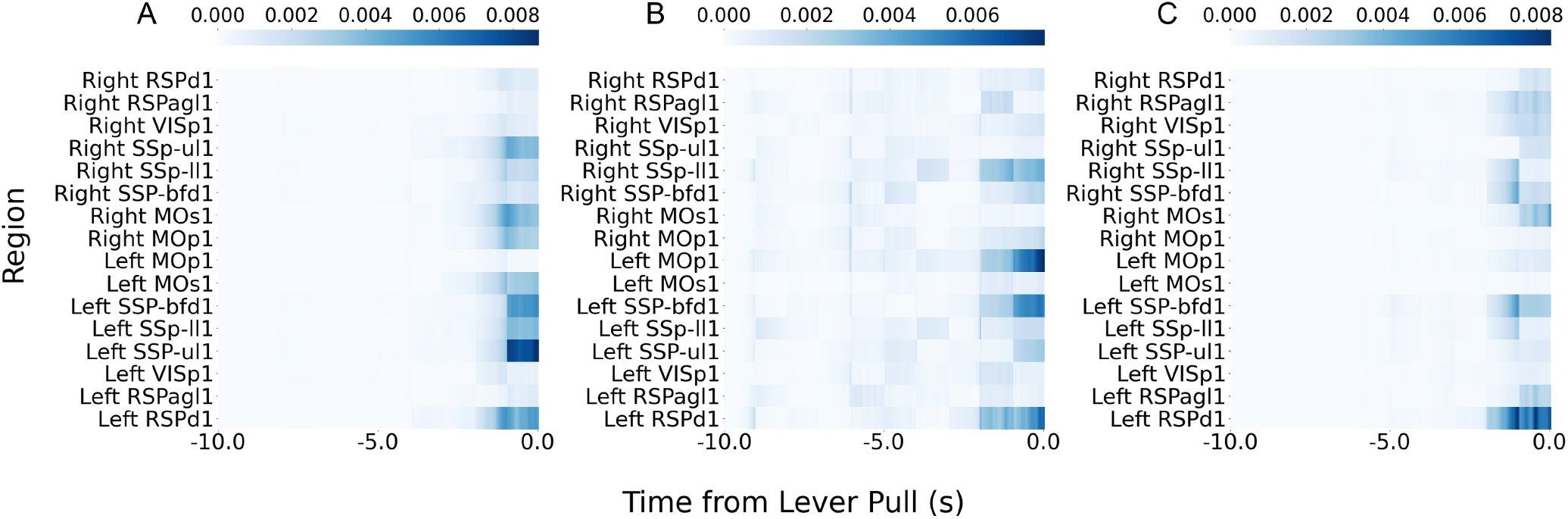
(A)(B)(C) Importance matrix of three example sessions with TV-RNN.

#### fMRI Data

We show the importance matrix of the fMRI dataset with TV-RNNs in Fig 11A,B. Here, the left pre-supplementary motor area (Left preSMA) is shown to be most important for both PD and healthy subjects. However, in healthy subjects, the difference of importance between Left preSMA and the other regions is not as much as the case of PD subjects. Additionally, all the most important regions for decoding are at the beginning of the sequence, which is different from the WFCI dataset.

**Figure 11.**
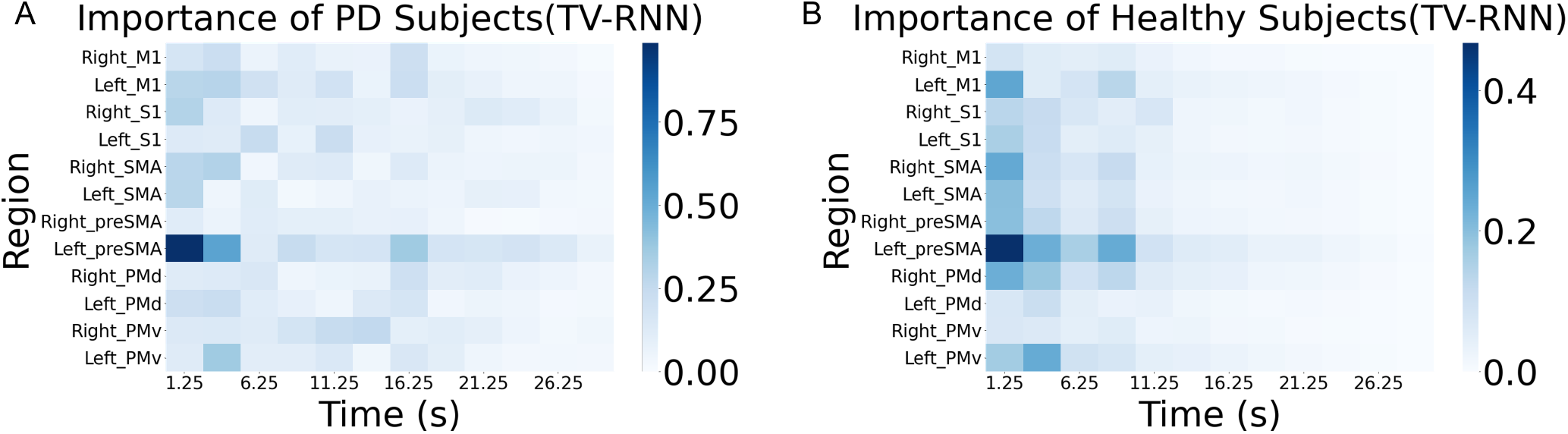
(A) Importance matrix of PD subjects with TV-RNNs; (B) Importance matrix of healthy subjects with TV-RNN.

## Conclusion

In this work, we developed a novel time varying model based on RNNs, to explore robust early sequential classification with brain-wide neural activity when the data distribution shifts across time. We show that TV-RNNs are able to achieve temporal robust classification earlier than standard RNNs and have higher accuracy. Moreover, we used the SHAP value to quantify the effect of different regions. We show somatosensory and motor areas at several seconds before the behavior are more important in behavioral decoding. The PreSMA region shows importance in both Parkinson’s patients and healthy controls. In the future, we aim at robust online decoding by using time series data, and using these models in the control theory setting to build adaptive controllers for neural data.

## Acknowledgements

We gratefully acknowledge support from the National Science Foundation grant 2219876 and National Institutes of Health grant (R01 NS058487, R01 NS052318).

## Supporting information

**Figure S1.**
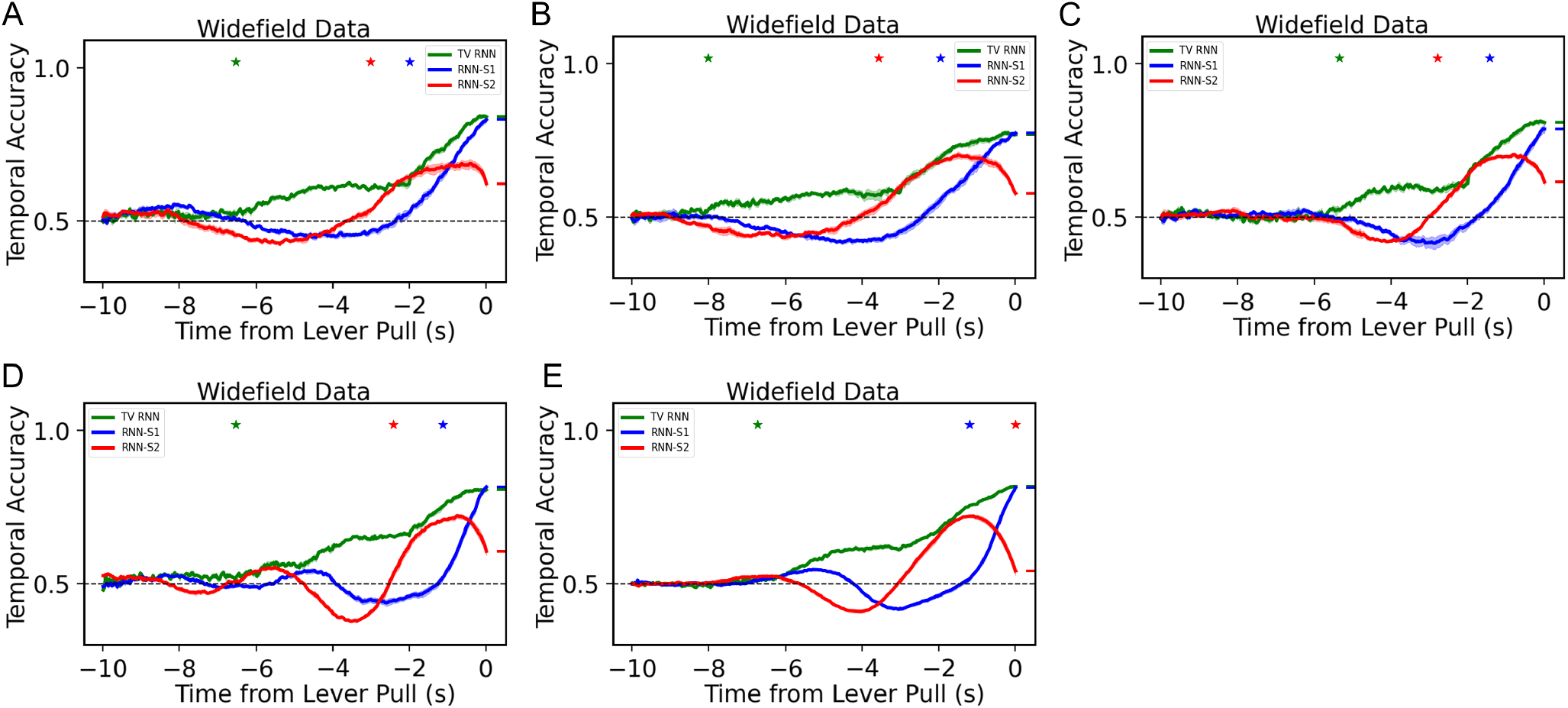
Temporal accuracy of standard RNNs with two training strategies and TV-RNNs for other 5 mice, the result of 1 mouse has shown in Fig 5B.

**Figure S2.**
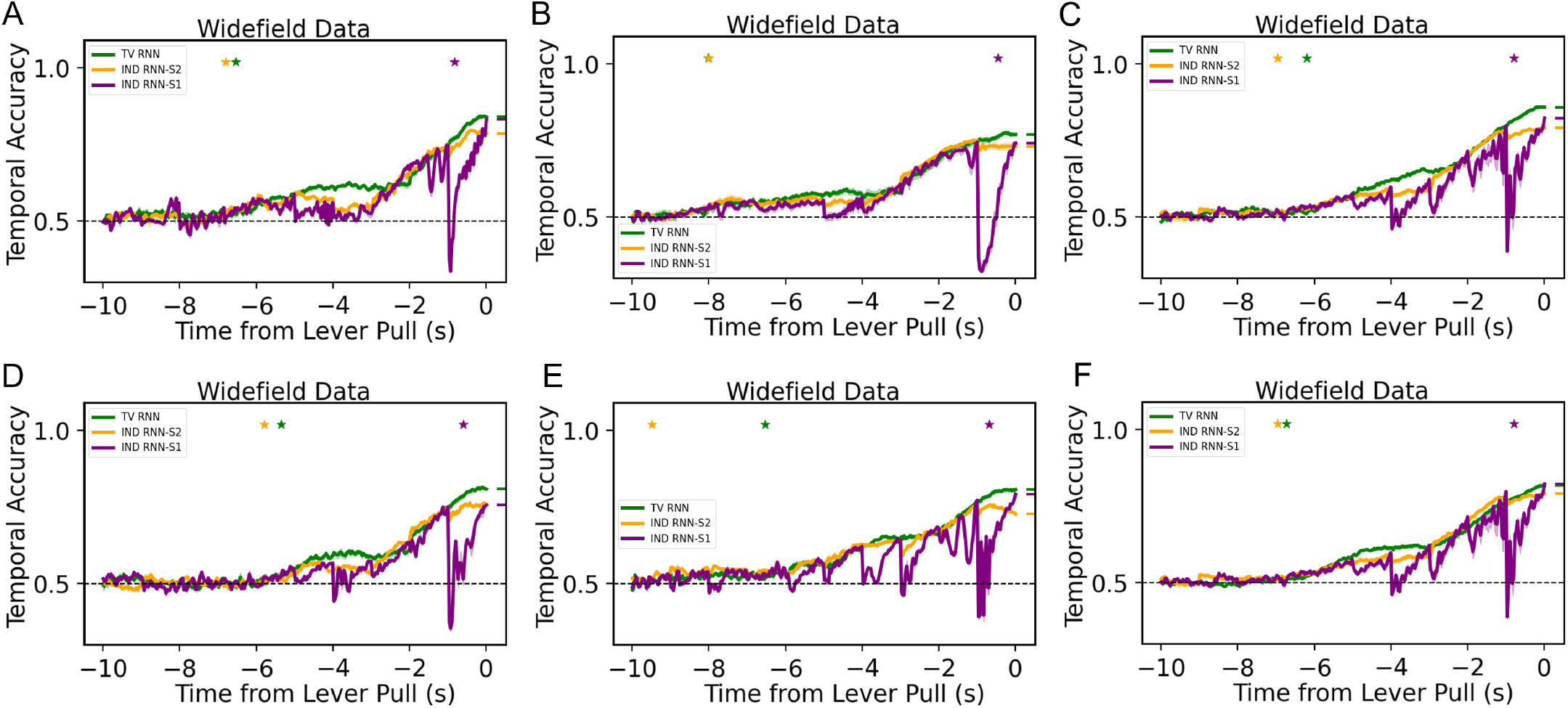
Temporal accuracy of independent standard RNNs with two training strategies and TV-RNNs for all 6 mice.

**Figure S3.**
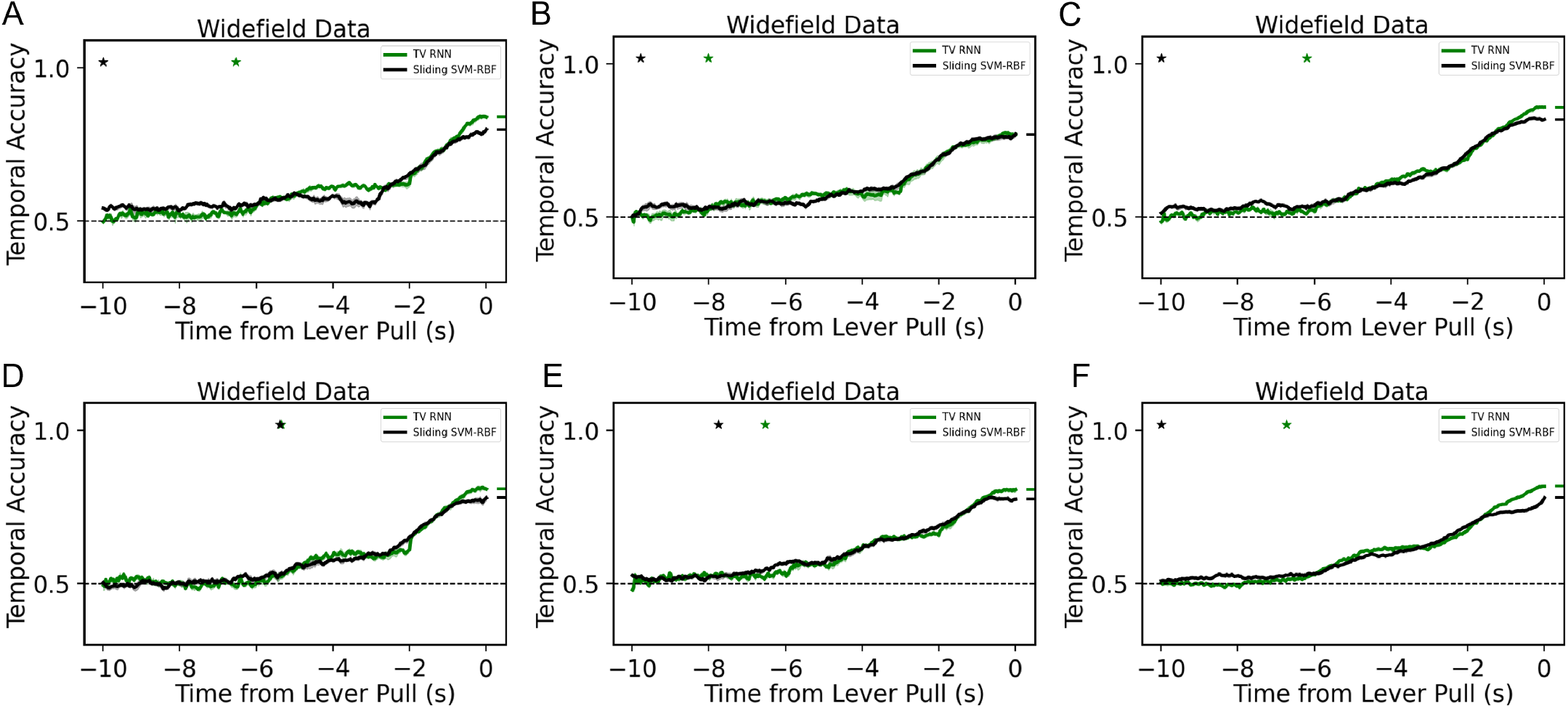
Temporal accuracy of sliding SVMs and TV-RNNs for all 6 mice.

**Figure S4.**
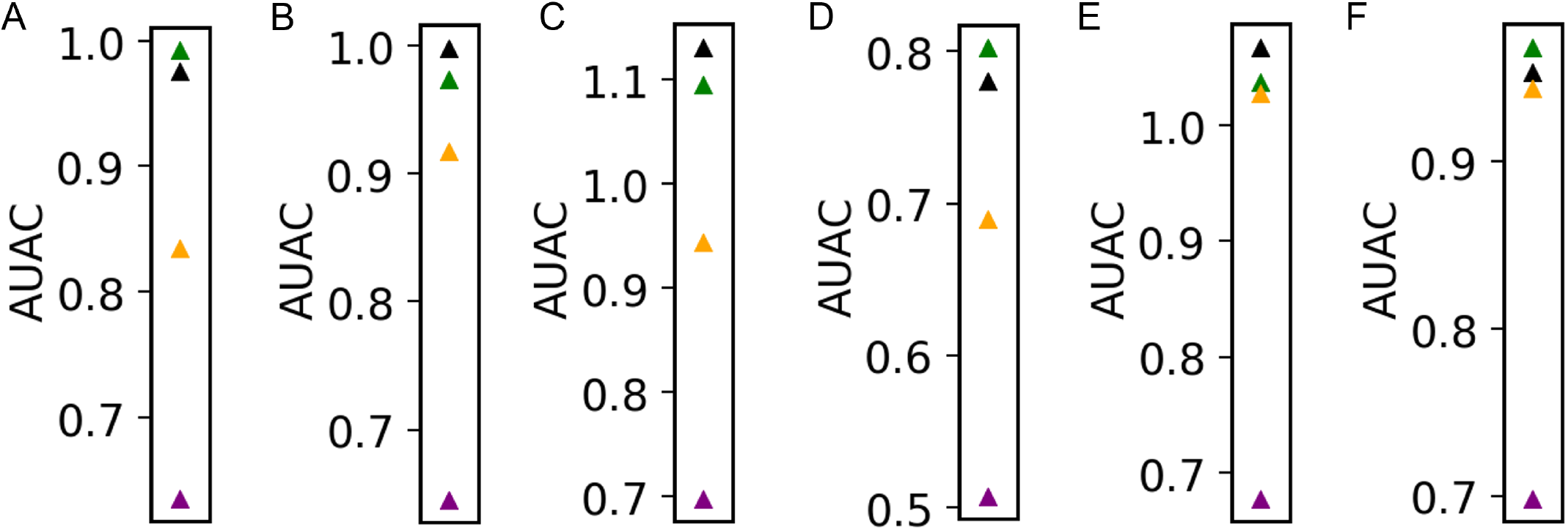
Area under accuracy curve using all classifiers described above for all 6 mice.

**Figure S5.**
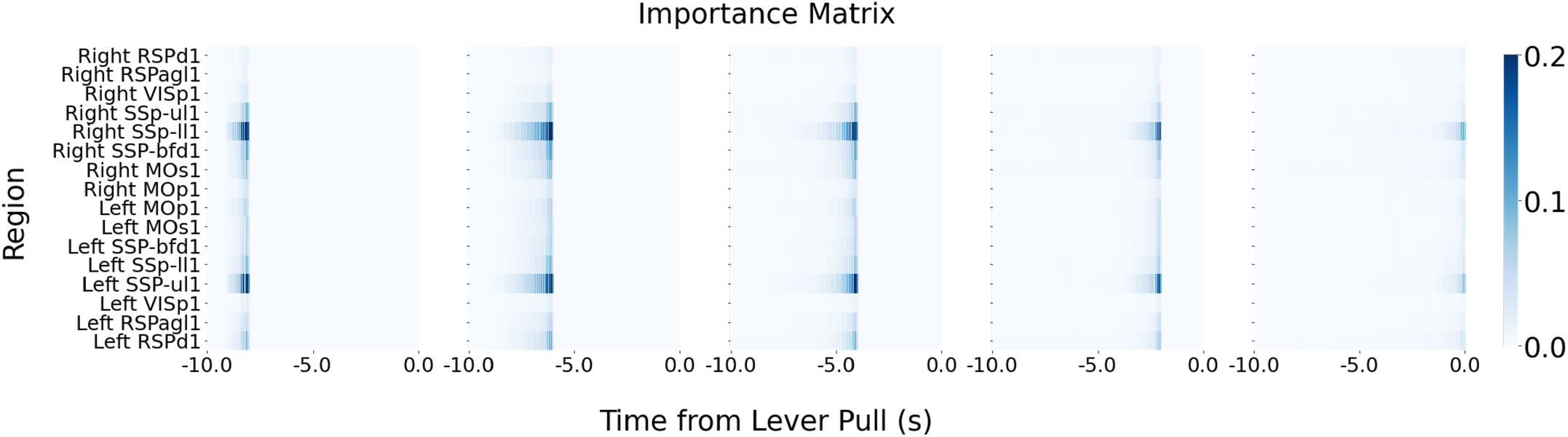
Importance matrix of one example mouse (trained with trials combined). The importance is from the mean absolute value of SHAP across trials by using different temporal output, i.e., from 8 seconds before the behavior to 0 second before the behavior.

